# Optimizing transcranial direct current stimulation (tDCS) electrode position, size, and distance doubles the on-target cortical electric field: Evidence from 3000 Human Connectome Project models

**DOI:** 10.1101/2021.11.21.469417

**Authors:** Kevin A. Caulfield, Mark S. George

## Abstract

Transcranial direct current stimulation (tDCS) is a widely used noninvasive brain stimulation technique with mixed results and no FDA-approved therapeutic indication to date. So far, thousands of published tDCS studies have placed large scalp electrodes directly over the intended brain target and delivered the same stimulation intensity to each person. Inconsistent therapeutic results may be due to insufficient cortical activation in some individuals and the inability to determine an optimal dose. Here, we computed 3000 MRI-based electric field models in 200 Human Connectome Project (HCP) participants, finding that the largely unexamined variables of electrode position, size, and between-electrode distance significantly impact the delivered cortical electric field magnitude. At the same scalp stimulation intensity, smaller electrodes surrounding the neural target deliver more than double the on-target cortical electric field while stimulating only a fraction of the off-target brain regions. This new optimized tDCS method can ensure sufficient cortical activation in each person and could produce larger and more consistent behavioral effects in every prospective research and transdiagnostic clinical application of tDCS.

## Introduction

Transcranial direct current stimulation (tDCS) is a widely used form of noninvasive brain stimulation(1) applied in healthy adults and over 20 neurological or psychiatric diagnoses, including depression(2–4), post-stroke motor rehabilitation(5, 6), aphasia(7–9), and anxiety(10, 11). By passing small electrical currents of 1 or 2mA through ‘anodal’ and ‘cathodal’ scalp electrodes, tDCS can open sodium-dependent ion channels in ‘underlying’ neurons(12), increasing intracellular calcium concentration(13, 14) and driving long term potentiation (LTP) effects(15). However, despite widespread interest in tDCS and appeal due to its noninvasive(16), inexpensive(17), and easily disseminatable nature(18–21), there are inconsistent effects of stimulation(22–26) and no FDA-approved indication for tDCS to date.

A possible explanation for the inconsistent tDCS results is that a suboptimal amount of stimulation is reaching the cortex at the intended target due in part to the same scalp dose being applied for each person(27–30). Converging lines of evidence support investigating whether higher amounts of stimulation might improve clinical tDCS effects. First, a cadaver study, which directly measured tDCS-induced cortical activity from implanted recording electrodes post-mortem, suggested that higher than 2mA stimulation doses are necessary to alter neuronal resting membrane potentials(27). Second, the emerging tool of electric field (Efield) modeling is an magnetic resonance imaging (MRI)-based method of accurately estimating how much stimulation reaches the cortex, and has shown that individuals receiving higher cortical E-fields from uniform 2mA dosing have greater therapeutic effects in depression(31) and working memory(32). Third, researchers have begun to test the safety and feasibility of higher tDCS intensities up to 4mA(28, 33, 34), with the goal of applying this in populations that may necessitate higher stimulation intensities such as in stroke patients. Taken together, there is building momentum behind using higher intensity tDCS to produce stronger and more consistent behavioral results, particularly in individuals who may have been underdosed at typical 2mA stimulation intensities. An alternative strategy put forth in this study is to develop more efficient tDCS techniques that use the same 2mA stimulation intensity at the scalp while producing larger E-fields at the cortex, effectively producing the same E-field at the cortex but possibly with fewer side effects (i.e. scalp burning) and less off-target stimulation inherent to higher stimulation intensities.

We show here, with extensive modeling in 200 Human Connectome Project (HCP) brains(35), that tDCS researchers have wrongly assumed where maximal stimulation effects are induced and thus have inefficiently stimulated the cortical target in nearly every application to date. At the cortical level, the maximal E-field is not underneath the electrodes, but rather between them. By using a novel electrode placement surrounding the intended cortical target with electrodes and by varying the electrode size, we demonstrate how to significantly increase the on-target electric field within the brain in comparison to conventional electrode placements, and with better accuracy (fewer off-target effects). In four rounds of E-field modeling, we demonstrate the effects of electrode position, size, and inter-electrode distance, and how optimizing these largely unexamined variables can more than double the on-target cortical Efield. In Round 1, we tested the effects of electrode <u>location</u>, and demonstrate how placing electrodes surrounding a cortical motor target influences the cortical E-field magnitude and focality of stimulation compared to conventional electrode montages in which electrodes are put directly over the cortical target. In Rounds 2 and 3, we systematically tested the effects of electrode <u>size</u> and inter-electrode <u>distance</u> on E-fields. Lastly, we optimized electrode placement, size, and distance in Round 4.

## Results

### Overview

We computed 3000 electric field models in 200 HCP participants, using the same 2mA stimulation intensity in each model and a within-subjects design (15 paired models per participant). Each model placed electrodes to target the motor cortex as a representative brain region commonly targeted in tDCS, with the idea that these principles could be translated to other cortical targets in the future. We used three outcome measures for each model (**Figure 1**): 1) E-field magnitude using a region of interest (ROI) analysis, with an individually placed 10mm radius spherical ROI in the primary motor cortex (M1) for each person. This ROI was kept constant across models, so any differences in ROI E-field were due to electrode position, size, or inter-electrode distance; 2) Whole brain maximum E-field magnitude using a 99^th^ percentile threshold to capture the average E-field of the top 1% of voxels irrespective of location; 3) Focality of electric fields by taking the volume (mm^3^) of electric fields equal to or greater than the 50^th^ percentile E-field. Lower volume stimulated indicated more focal stimulation.

**Figure 1:**
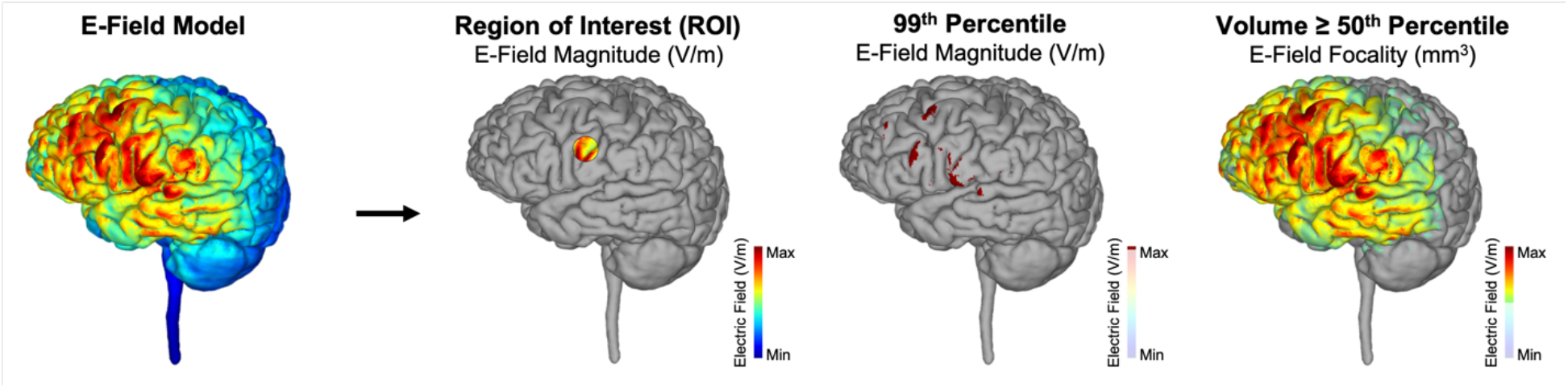
Electric Field Modeling Outcome Measures. For each E-field model, we performed three types of analyses: 1) Region of Interest (ROI) E-Field Magnitude. This method examines the E-field magnitude in a 10mm radius spherical ROI that was individually placed underneath the cortical projection at C3. This ROI was the same for every model and represented the Efield at the on-target cortical motor target. 2) 99^th^ Percentile E-Field Magnitude. This method determined the E-field magnitude at the whole brain level, examining the top 1% of activated voxels irrespective of whether the effects were on-target or off-target. 3) E-Field Focality. We measured focality by measuring the volume of voxels (mm^3^) equal to or above the 50^th^ percentile E-field. The greater the volume stimulated, the lower the focality of stimulation (greater off-target effects).

### Round 1: Effects of Electrode Location in Conventional vs. Novel Surround Electrode Montages

In the first five montages, we tested the effects of electrode location on the E-field magnitude and focality in three conventional electrode montages (bilateral M1, M1-supraorbital (SO), and high definition (HD)-tDCS)(36, 37) and two novel placements surrounding the cortical target (left right pad surround (LRPS)-tDCS and anterior posterior pad surround (APPS)-tDCS)(**Figure 2a**). LRPS and APPS montages placed electrodes equidistant from the motor cortical target using electroencephalography (EEG) coordinates (LRPS: anode at C1, cathode at C5 and APPS: anode at CP3, cathode at FP3)(**Figure 2a**) with the goal of centering the maximum E-field at the cortical motor target midway between electrodes. All pad electrodes had 7 × 5cm dimensions while HD-tDCS had 0.5cm diameter circular electrodes, with the HD-tDCS anode at C3 and HD-tDCS cathodes at FC3, C1, CP3, and C5. We projected the results of all 200 models (1 per person, paired between models) per montage into average space in axial and sagittal 3D orientations in **Figure 2b-c**.

**Figure 2:**
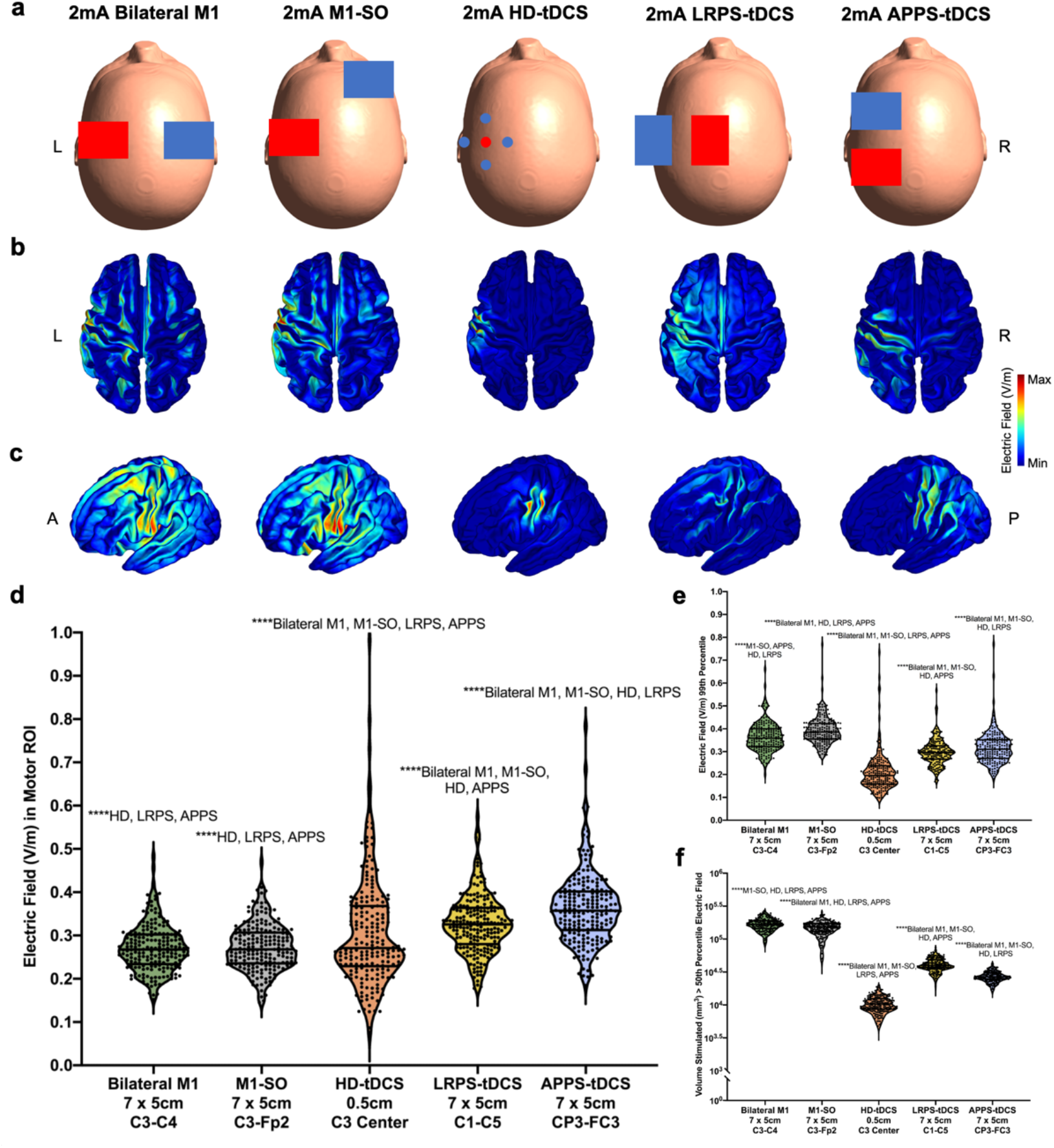
Round 1 Modeling: Electrodes Surrounding the Intended Cortical Target Produce Higher and More Focal Electric Fields. **a) Electrode Placements in Bilateral M1, M1-SO, HD-tDCS, LRPS-tDCS, and APPS-tDCS Montages**. Red = anodal electrodes, blue = cathodal electrodes. **b-c) E-Fields from 200 Models Projected Into fsaverage Space for Axial (b) and Sagittal (c) Orientations**. In each projection, 200 individual models were averaged into fsaverage space to visualize the E-field magnitude and spread for each electrode orientation. Notably, in conventional electrode montages, the maximal E-fields are not under the electrodes whereas in LRPS and APPS-tDCS the main effects are under the electrodes and focused at the intended target. **d) E-Field Magnitude in Motor ROI**. Electrodes surrounding the intended motor cortical target produced significantly higher E-fields in the ROI (****p < 0.0001). In particular, APPS-tDCS produced a higher E-field of 0.363V/m than in every other electrode placement. See **Table 1** for details. **e) E-Field Magnitude Using the 99^th^ Percentile Threshold**. The ranking of 99^th^ percentile E-field magnitudes differed from the ROI measurement, with conventional bilateral M1 and M1-SO montages producing the largest Efields. This suggests that characteristics of the bilateral M1 and M1-SO montages could be utilized to further optimize APPS-tDCS. **f) E-Field Focality**. HD-tDCS produced the most focal E-fields, with the lowest volume equal to or above the 50^th^ percentile E-field. This was followed by the two novel electrode placements, APPS-tDCS and LRPS-tDCS. The two conventional electrode placements of bilateral M1 and M1-SO produced the lowest focality E-fields. See **Table 1** for details.

The various electrode montages produced significantly different electric fields at the motor ROI, F(1.8, 361.4) = 169.0, p < 0.0001. Post-hoc analyses with Tukey corrections indicated that each comparison statistically differed at the p < 0.0001 significance level except for the conventional bilateral M1 and M1-SO montages. In particular, APPS-tDCS electrodes that surround the cortical motor target by placing electrodes in the anterior and posterior directions produced the highest mean E-field of 0.363V/m (SD = 0.081V/m), compared to LRPS-tDCS (mean = 0.326V/m, SD = 0.067V/m), HD-tDCS (mean = 0.306V/m, SD = 0.126V/m), bilateral M1 (mean = 0.272V/m, SD = 0.054V/m), and M1-SO (mean = 0.273V/m, SD = 0.053V/m)(**Figure 2d**; **Table 1**).

**Table 1:**
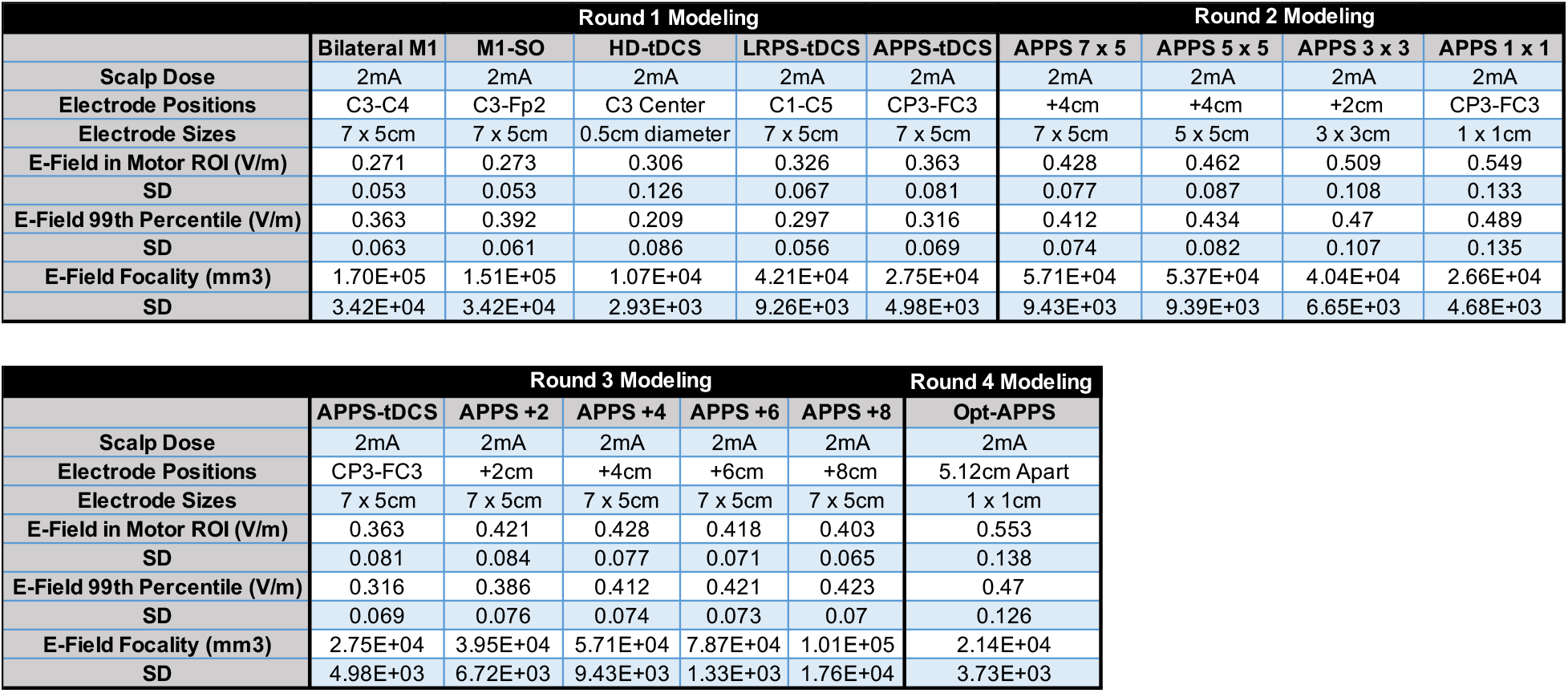
Means and Standard Deviations of E-Field Magnitudes and Focality in Rounds 1-4 Modeling.

|  | Round 1 Modeling |  |  |  |  | Round 2 Modeling |  |  |  |
| --- | --- | --- | --- | --- | --- | --- | --- | --- | --- |
|  | Bilateral M1 | M1-SO | HD-tDCS | LRPS-tDCS | APPS-tDCS | APPS 7 x 5 | APPS 5 x 5 | APPS 3 x 3 | APPS 1 x 1 |
| Scalp Dose | 2mA | 2mA | 2mA | 2mA | 2mA | 2mA | 2mA | 2mA | 2mA |
| Electrode Positions | C3-C4 | C3-Fp2 | C3 Center | C1-C5 | CP3-FC3 | +4cm | +4cm | +2cm | CP3-FC3 |
| Electrode Sizes | 7 x 5cm | 7 x 5cm | 0.5cm diameter | 7 x 5cm | 7 x 5cm | 7 x 5cm | 5 x 5cm | 3 x 3cm | 1 x 1cm |
| E-Field in Motor ROI (V/m) | 0.271 | 0.273 | 0.306 | 0.326 | 0.363 | 0.428 | 0.462 | 0.509 | 0.549 |
| SD | 0.053 | 0.053 | 0.126 | 0.067 | 0.081 | 0.077 | 0.087 | 0.108 | 0.133 |
| E-Field 99th Percentile (V/m) | 0.363 | 0.392 | 0.209 | 0.297 | 0.316 | 0.412 | 0.434 | 0.47 | 0.489 |
| SD | 0.063 | 0.061 | 0.086 | 0.056 | 0.069 | 0.074 | 0.082 | 0.107 | 0.135 |
| E-Field Focality (mm3) | 1.70E+05 | 1.51E+05 | 1.07E+04 | 4.21E+04 | 2.75E+04 | 5.71E+04 | 5.37E+04 | 4.04E+04 | 2.66E+04 |
| SD | 3.42E+04 | 3.42E+04 | 2.93E+03 | 9.26E+03 | 4.98E+03 | 9.43E+03 | 9.39E+03 | 6.65E+03 | 4.68E+03 |

|  | Round 3 Modeling |  |  |  |  | Round 4 Modeling |
| --- | --- | --- | --- | --- | --- | --- |
|  | APPS-tDCS | APPS +2 | APPS +4 | APPS +6 | APPS +8 | Opt-APPS |
| Scalp Dose | 2mA | 2mA | 2mA | 2mA | 2mA | 2mA |
| Electrode Positions | CP3-FC3 | +2cm | +4cm | +6cm | +8cm | 5.12cm Apart |
| Electrode Sizes | 7 x 5cm | 7 x 5cm | 7 x 5cm | 7 x 5cm | 7 x 5cm | 1 x 1cm |
| E-Field in Motor ROI (V/m) | 0.363 | 0.421 | 0.428 | 0.418 | 0.403 | 0.553 |
| SD | 0.081 | 0.084 | 0.077 | 0.071 | 0.065 | 0.138 |
| E-Field 99th Percentile (V/m) | 0.316 | 0.386 | 0.412 | 0.421 | 0.423 | 0.47 |
| SD | 0.069 | 0.076 | 0.074 | 0.073 | 0.07 | 0.126 |
| E-Field Focality (mm3) | 2.75E+04 | 3.95E+04 | 5.71E+04 | 7.87E+04 | 1.01E+05 | 2.14E+04 |
| SD | 4.98E+03 | 6.72E+03 | 9.43E+03 | 1.33E+03 | 1.76E+04 | 3.73E+03 |

The 99^th^ percentile F-test was also significant, F(1.9, 371.9) = 843.6, p < 0.0001 (**Figure 2e; Table 1**). Post-hoc Tukey corrected tests revealed that the conventional bilateral M1 and M1-SO montages produced the two highest 99^th^ percentile E-fields of 0.363V/m (SD = 0.063V/m) and 0.392V/m (SD = 0.061V/m) respectively. These values were significantly higher than HD-tDCS (mean = 0.209V/m, SD = 0.086V/m), LRPS-tDCS (mean = 0.297V/m, SD = 0.056V/m), and APPS-tDCS (mean = 0.316V/m, SD = 0.069V/m). Each comparison was significant at the p < 0.0001 significance level.

The omnibus test for focality showed significantly different spreads of electric fields between electrode placements, F(2.0, 402.8) = 2975, p < 0.0001 (**Figure 2f; Table 1**). Tukey corrected post-hoc comparisons revealed that the volume stimulated from each electrode montage significantly differed at the p < 0.0001 significance level. HD-tDCS delivered the most focal amount of stimulation (lowest volume stimulated, fewest off-target effects) at 1.07 × 10^4^ mm^3^ (SD = 2.93 × 10^3^ mm^3^). Of the pad electrode montages, APPS-tDCS delivered the most focal amount of stimulation (mean = 2.75 × 10^4^ mm^3^, SD = 4.98 × 10^3^ mm^3^), followed by LRPS-tDCS (mean = 4.21 × 10^4^ mm^3^, SD = 9.26 × 10^3^ mm^3^), M1-SO (mean = 1.51 × 10^5^ mm^3^, SD = 3.42 × 10^4^ mm^3^), and bilateral M1 (mean = 1.70 × 10^5^ mm^3^, SD = 3.42 × 10^4^ mm^3^). In sum, the APPS-tDCS paradigm produced the highest motor ROI E-field and with the greatest focality amongst non-HD-tDCS montages.

### Location of the Maximal Electric Field Underneath vs. Between APPS-tDCS Electrodes

Round 1 of modeling showed that the maximal E-field is between and not underneath the electrodes since positioning the electrodes surrounding the cortical motor target produced higher E-fields in the motor ROI. As a second method of investigating the location of the maximal E-field, and to confirm that this effect is not only observed at a group level but also at an individual level, we examined the E-field magnitude at three ROIs for the APPS-tDCS 7 × 5cm model. We placed the three ROIs at the same motor cortical target underneath C3, and at individually placed cortical grey matter projections underneath the center of the anodal (CP3) and cathodal (FC3) electrodes using the same 10mm radius spherical ROI shape (**Figure 3a-b**). Quantitatively (**Figure 3c**) and visually (**Figure 3d**), the group level maximal E-field was midway between electrodes at the motor ROI (mean = 0.363V/m, SD = 0.081V/m), as opposed to underneath the anode at 0.172V/m (SD = 0.043V/m) and cathode at 0.201V/m (SD = 0.052V/m). Furthermore, this effect was consistent between individuals, with 200 of 200 participants having the maximal E-field midway between, and not underneath, the electrodes. Thus, the maximal E-field is repeatedly induced midway between the electrodes in each person, further affirming the utility in using APPS-tDCS to place electrodes surrounding the cortical target to maximize the E-field.

**Figure 3:**
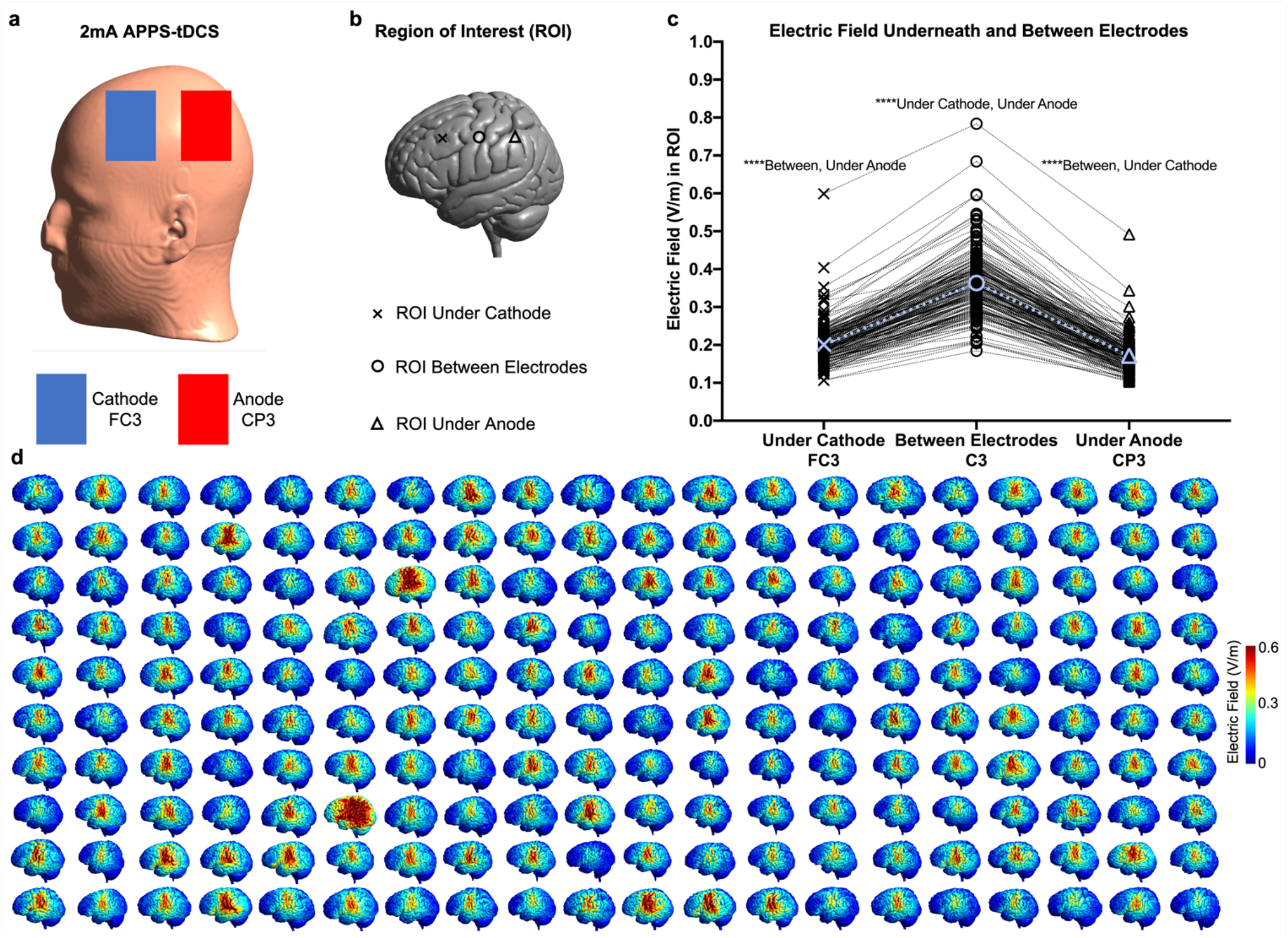
tDCS Induces Maximal Electric Fields Between, Not Underneath Electrodes. **a) Placement of Anodal (Red) and Cathodal (Blue) Electrodes in Each Individual Head Model. b) Location of Regions of Interest (ROIs).** We measured the E-field at a 10mm radius spherical ROI at the grey matter location immediately underneath the center of the anodal and cathodal electrodes, and at the same motor cortical ROI used in each round of modeling (cortical projection underneath the C3 electrode location). This allowed us to examine the difference between the E-fields induced underneath vs. between each electrode in the APPS-tDCS montage. **c) Electric Field Underneath and Between Electrodes**. The maximum E-field for each of the 200 participants was midway between the electrodes (****p < 0.0001), affirming the APPS-tDCS strategy of surrounding the cortical target for maximizing induced E-fields. The E-field between each electrode (mean = 0.363V/m) was significantly greater than that of the Efields underneath the anode (mean = 0.172V/m) and cathode (mean = 0.201V/m). Averages are denoted in blue with dashed lines. **d) Visualization of APPS-tDCS Modeling (N = 200**). Qualitatively, APPS-tDCS induced the maximum E-field midway between, not underneath, electrodes in 200 individual head models. This visualization substantiates that the maximal Efield is consistently induced midway between tDCS electrodes.

### Similarity Analyses Between ROI Motor Target and 99^th^ Percentile Whole Brain Peak Electric Fields

A third method to assess whether a given electrode montage induces the peak electric field at the intended cortical target is to compare the similarity between the whole brain 99^th^ percentile and region specific ROI electric fields. If the E-field in the top 1% of voxels were higher than the ROI electric field, the tDCS montage delivers the maximal E-field outside of the cortical target. Alternatively, if the ROI electric field were higher than or equivalent to the 99^th^ percentile electric field, the cortical target is located within the top 1% of activated voxels. For each participant, we plotted the motor ROI E-field (blue) and 99^th^ percentile E-field (red) and computed an intraclass correlation coefficient (ICC) as a statistical measure of similarity between the ROI and 99^th^ percentile E-fields in the five Round 1 models (bilateral M1, M1-SO, HD-tDCS, LRPS-tDCS, and APPS-tDCS)(**Figure 4a-e**).

**Figure 4:**
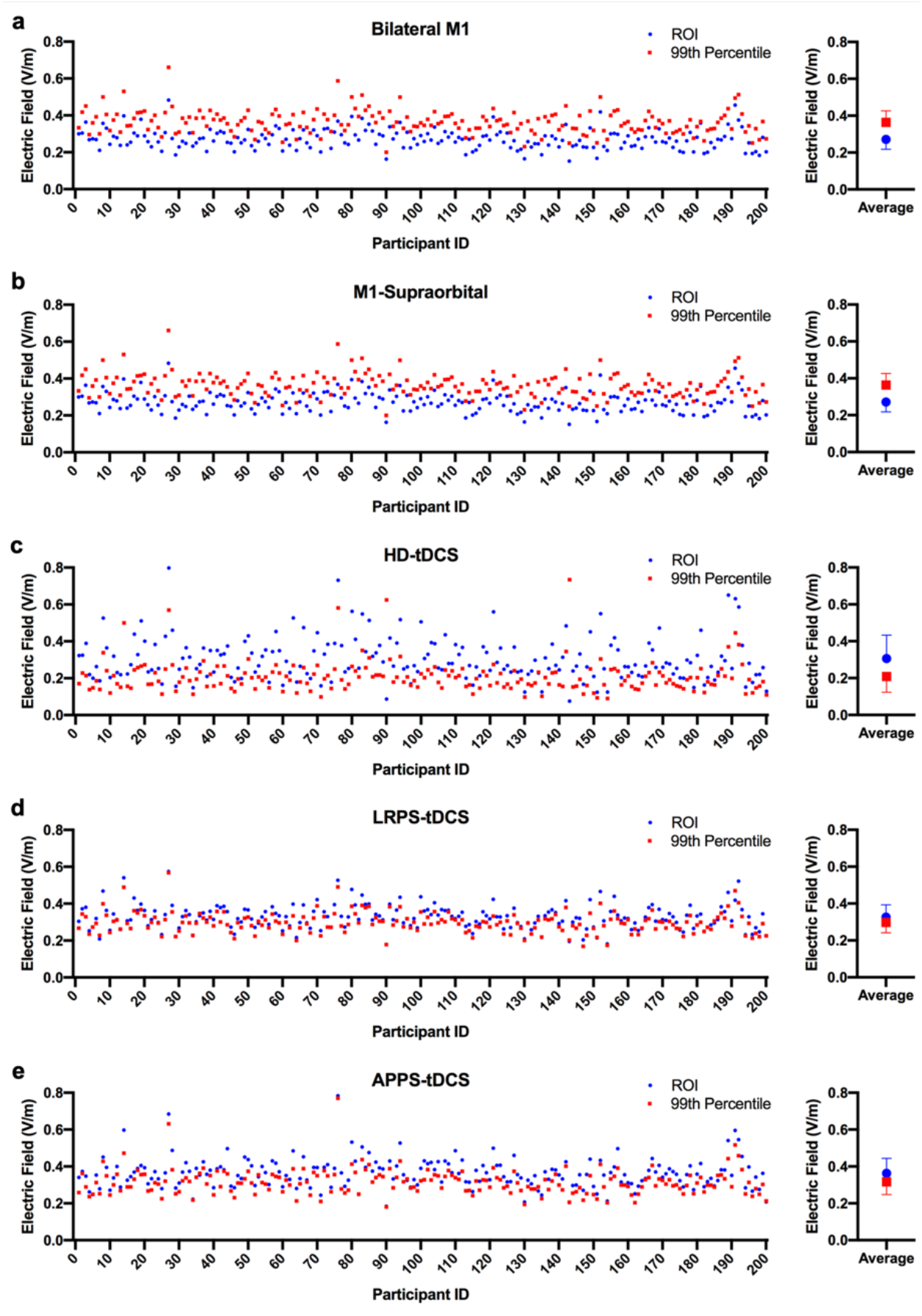
APPS-tDCS Induces the Maximal E-Field at the Intended Cortical Target While Conventional Electrode Placements Focus Stimulation Off-Target. In this figure, we compared E-field magnitudes between the motor target (as assessed by the region of interest; ROI analysis), and the whole brain peak E-fields using the 99^th^ percentile threshold (top 1% of voxels included). If these measures are relatively similar, as assessed by an intraclass correlation coefficient (ICC), the whole brain peak E-field is situated at the cortical target ROI. If the 99^th^ percentile E-field exceeds that of the ROI, the maximal E-field is located outside of the cortical target. All ICCs were significant at the p < 0.0001 level, but with differing levels of consistency. For Bilateral M1 (**a**) and M1-SO (**b**), the 99^th^ percentile E-field consistently exceeded that of the ROI in each of the 200 participants, with ICCs of 0.391 and 0.241 respectively (‘poor’ reliability). For HD-tDCS (**c**) the ROI E-field was consistently higher than the 99^th^ percentile E-field, suggesting that HD-tDCS stimulated a cortical target with a very focal spread of stimulation, as the ROI E-field was in a more constrained space than that of the top 1% of voxels, but with a ‘poor’ reliability ICC of 0.430. For LRPS-tDCS (**d**) and APPS-tDCS (**e**), there was ‘good’ similarity between the ROI and 99^th^ percentile E-fields of 0.831 and 0.772 respectively, suggesting that the maximal E-field was located within the cortical target ROI in each of the 200 participants. These ICC analyses provide further evidence that placing electrode surrounding the cortical target, as we did in LRPS and APPS-tDCS, places the maximal E-field at the target region.

The ICC results consistently demonstrated that conventional electrode placements deliver maximal E-fields off-target whereas surrounding the intended target focuses the highest E-field within the motor cortex. For Bilateral M1 (**Figure 4a**) and M1-SO tDCS (**Figure 4b**), the 99^th^ percentile E-field was higher than the ROI E-field in 200 of 200 participants, suggesting that these conventional electrode placements consistently do not deliver the maximal E-field at the cortical motor target. While each comparison was statistically significant at the p < 0.001 significance level, the bilateral M1 ICC of 0.391 and M1-Supraorbital ICC of 0.241 fall within the ‘poor’ reliability range. HD-tDCS had higher ROI than 99^th^ percentile E-field values in 200 of 200 participants due to the high focality of stimulation, with a ‘poor’ reliability ICC value of 0.430 (**Figure 4e**). In contrast, for APPS-tDCS and LRPS-tDCS in which the electrodes surround the cortical target, there were statistically significant and ‘good’ similarity values between the 99^th^ percentile and ROI E-fields (APPS-tDCS ICC: 0.772; LRPS-tDCS ICC: 0.831), with the motor ROI values being higher than the whole brain 99^th^ percentile values in 200 of 200 participants for both electrode placements (**Figure 4c-d**). Thus, these novel surround electrode montages consistently deliver maximal E-fields on-target at the intended cortical target; the conventional placements of bilateral M1 and M1-SO focus the maximal amount of stimulation consistently off-target and therefore are inefficient methods of stimulating the intended cortical target region.

### Round 2: Isolating the Effect of Electrode Size on Electric Field Magnitude and Focality

After showing the utility of novel tDCS surround electrode montages in Round 1 modeling, we examined ways of further optimizing APPS-tDCS. In **Supplementary Section 1**, we examined the effects of electrode size and distance on E-field magnitude and focality, but unintentionally conflated these two variables. This spurious approach centered each electrode of different sizes at the same CP3-FC3 coordinates, unintentionally causing the edge-to-edge inter-electrode distance varied between an average of 3.12cm in the 7 × 5cm montage to 7.12cm in the 1 × 1cm montage. In order to disambiguate these variables, Rounds 2 and 3 of modeling systematically isolated the effects of electrode size and distance on E-field magnitude and focality.

In Round 2 of modeling, we placed the 1 × 1cm electrodes at the CP3-FC3 position for each participant (average of 7.12cm apart). We then individually matched the edge-to-edge distance between electrodes by placing 3 × 3cm, 5 × 5cm, and 7 × 5cm electrodes the same mean 7.12cm distance apart to isolate the variable of electrode size on electric field magnitude and focality (**Figure 5a-c**).

**Figure 5:**
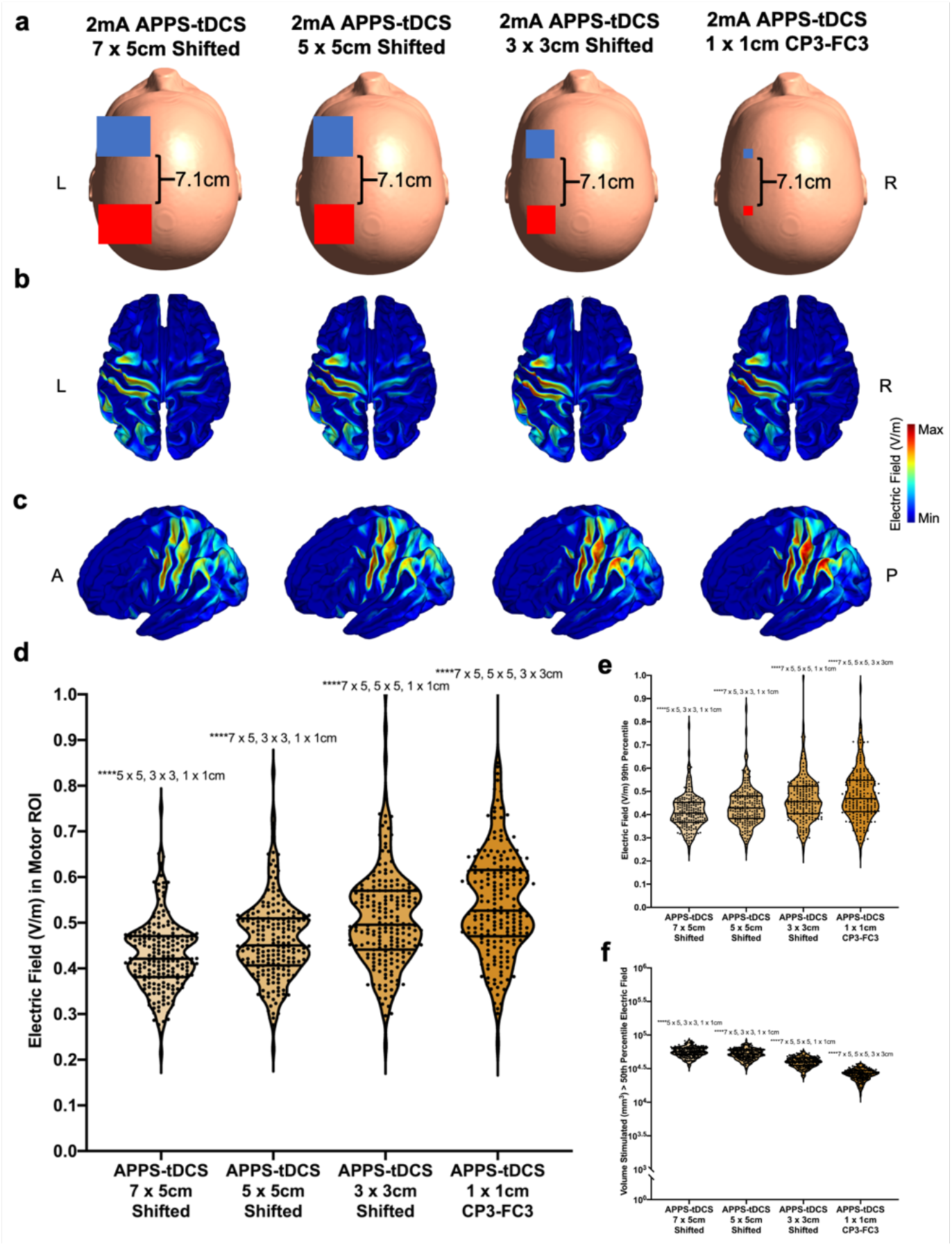
Round 2 Modeling: Smaller Electrodes Produce Significantly Higher and More Focal Electric Fields at the Intended Cortical Target. **a) Electrode Sizes and Positioning.** In order to isolate the effects of electrode size on E-field magnitude and focality, we matched the edge-to-edge distance between electrodes Red = anode, blue = cathode. **b-c) E-Fields from 200 Models Projected Into fsaverage Space for Axial (b) and Sagittal (c) Orientations**. Visually, the E-fields from the smaller 1 × 1cm electrodes were higher than those produced by the 3 × 3cm, 5 × 5cm, and 7 × 5cm sizes. **d) E-Field Magnitude in Motor ROI**. APPS-tDCS with 1 × 1cm electrodes produced the highest E-field of 0.549V/m, compared to 3 × 3cm (mean = 0.509V/m), 5 × 5cm (mean = 0.462V/m), and 7 × 5cm electrodes (mean = 0.428V/m). **e) E-Field Magnitude Using the 99^th^ Percentile Threshold**. 99^th^ percentile E-fields followed a similar pattern as the motor ROI E-fields, with 1 × 1cm electrodes producing the highest magnitude E-fields (all comparisons ****p < 0.0001; see **Table 1** for details). **f) E-Field Focality**. 1 × 1cm electrodes stimulated the most focal volume, with smaller electrode sizes being inversely related to spread of E-fields (****p < 0.0001). In sum, electrode size inversely relates to the E-field magnitude such that smaller electrodes produce higher and more focal Efields (fewer off-target effects).

The electric field magnitude significantly differed by electrode size in both the motor ROI target (F(1.1, 209.9) = 730.4, p < 0.0001) and with the whole brain 99^th^ percentile threshold (F(1.0, 208.7) = 236.5, p < 0.0001)(**Figure 5d-e; Table 1**). These values inversely scaled as a function of electrode size, such that the smallest 1 × 1cm electrode size had the highest E-field at the motor ROI (mean = 0.549V/m, SD = 0.133V/m), compared to 3 × 3cm (mean = 0.509V/m, SD = 0.108V/m), 5 × 5cm (mean = 0.462V/m, SD = 0.089V/m), and 7 × 5cm (mean = 0.428V/m, SD = 0.077V/m), with each comparison being statistically significant at the Tukey-corrected p < 0.0001 level (**Figure 5d**). Similarly, the 99^th^ percentile approach yielded similar results, all at the p < 0.0001 significance level, with the 1 × 1cm electrodes again producing the highest whole brain peak E-field of 0.489V/m (SD = 0.135V/m), followed by the 3 × 3cm (mean = 0.47V/m, SD = 0.107V/m), 5 × 5cm (mean = 0.434V/m, SD = 0.082V/m), and 7 × 5cm electrodes (mean = 0.412V/m, SD = 0.074V/m)(**Figure 5e**).

Lastly, focality also inversely scaled as a function of electrode size, such that the smallest electrodes produced the most focal E-field, F(1.3, 251.4) = 3398, p < 0.0001 (**Figure 5f; Table 1**). The 1 × 1cm electrodes stimulated the smallest volume ≥ 50^th^ percentile E-field of 2.66 × 10^4^ mm^3^ (SD = 4.68 × 10^3^ mm^3^), compared to 3 × 3cm (mean = 4.04 × 10^4^ mm^3^, SD = 6.65 × 10^3^ mm^3^), 5 × 5cm (mean = 5.37 × 10^4^ mm^3^, SD = 9.39 × 10^3^ mm^3^), and 7 × 5cm electrodes (mean = 5.71 × 10^4^ mm^3^, SD = 9.43 × 10^3^ mm^3^).

### Round 3: Isolating the Effect of Electrode Distance on Electric Field Magnitude and Focality

Round 3 of modeling kept electrode size constant at the 7 × 5cm size and moved electrodes farther apart by +2cm increments to evaluate the impact of electrode distance on electric field magnitude and focality. This altered the edge-to-edge distance between electrodes from an average of 3.12cm apart at the original CP3-FC3 positioning, up to 11.12cm apart (**Figure 6a-c**).

**Figure 6:**
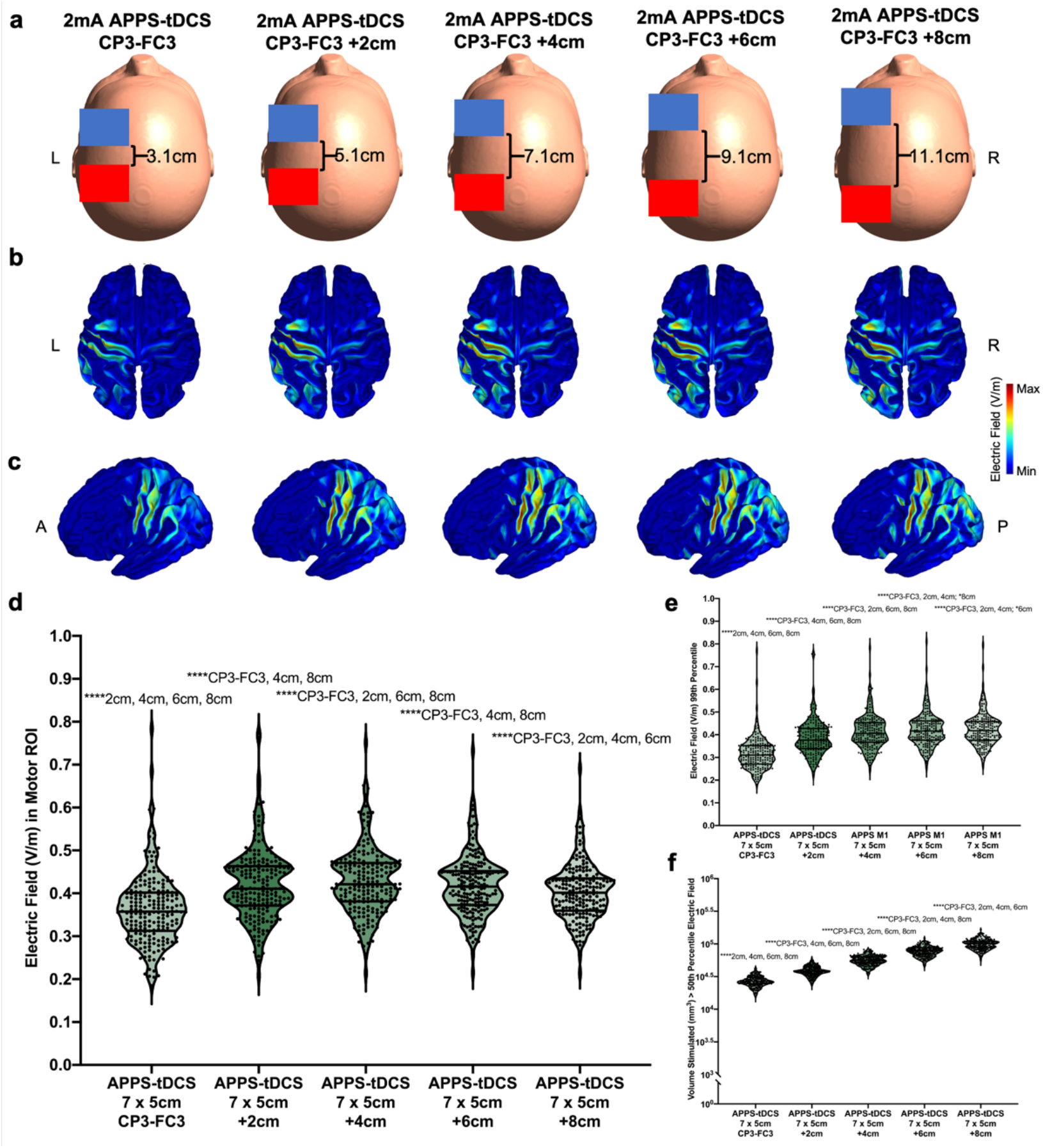
Round 3 Modeling: Inter-Electrode Distance Impacts Electric Field Magnitude and Focality of Stimulation. **a) Electrode Sizes and Positioning.** We kept the electrode size constant at 7 × 5cm and altered the positioning from the initial, individually placed electrode center locations of CP3-FC3 in +2cm increments up to +8cm (mean inter-electrode distance = 11.12cm). Red = anode, blue = cathode. **b-c) E-Fields from 200 Models Projected Into fsaverage Space for Axial (b) and Sagittal (c) Orientations**. Qualitatively, the maximal E-field increased from the initial CP3-FC3 placement to +2cm and +4cm. **d) E-Field Magnitude in Motor ROI**. All cortical target ROI E-fields significantly differed, such that the +2cm (mean = 0.421V/m) and +4cm (mean = 0.428V/m) produced higher E-fields than the CP3-FC3, +6cm, and +8cm placements (****p < 0.0001). **e) E-Field Magnitude Using the 99^th^ Percentile Threshold**. 99^th^ percentile E-fields linearly scaled with distance apart; larger inter-electrode distance corresponded with higher global E-fields (****p < 0.0001). **f) E-Field Focality**. The amount of volume stimulated linearly increased with distance apart, such that the APPS-tDCS at +8cm apart from the original placement was least focal, with the most off-target effects (****p < 0.0001). Round 3 modeling revealed that larger distance, to an extent, can facilitate higher Efields at the cortical target. However, the researcher also must take focality into consideration, as the +2cm and +4cm distances that induced higher cortical target E-fields additionally stimulated more volume.

Electrode distance significantly affected E-field magnitude in both the motor ROI, F(1.4, 274) = 502.8, p < 0.0001, and 99^th^ percentile threshold, F(1.4, 273.7) = 2065, p < 0.0001 (**Figure 6d-e; Table 1**). In the motor ROI, there was a significant increase in E-field magnitude from the initial APPS-tDCS model with 7 × 5cm electrodes placed at CP3-FC3 (mean = 0.363V/m, SD = 0.081V/m) to +2cm (mean = 0.428V/m, SD = 0.077V/m), +4cm (mean = 0.421V/m, SD = 0.084V/m), +6cm (mean = 0.418V/m, SD = 0.071V/m), and +8cm (mean = 0.403V/m, SD = 0.065V/m). Each comparison of E-field magnitude at the motor ROI significantly differed at the p < 0.0001 significance level except for +2 vs. +6cm. Similarly, the 99^th^ percentile E-field significantly differed (p < 0.0001) for each comparison and increased from the initial model (mean = 0.316V/m, SD = 0.069V/m) to +2cm (mean = 0.386V/m, SD = 0.076V/m), +4cm (mean = 0.412V/m, SD = 0.074V/m), +6cm (mean = 0.421V/m. SD = 0.073V/m), and +8cm (mean = 0.423V/m, SD = 0.07V/m).

For focality, the inter-electrode distance linearly affected the amount of volume stimulated, F(1.2, 243.1) = 3821, p < 0.0001 (**Figure 6f; Table 1**). The baseline CP3-FC3 APPS-tDCS positioning stimulated a volume of 2.75 × 10^4^ mm^3^ (SD = 4.98 × 10^3^ mm^3^), +2cm stimulated 3.96 × 10^4^ mm^3^ (SD = 6.72 × 10^3^ mm^3^), +4cm stimulated 5.71 × 10^4^ mm^3^ (SD = 9.43 × 10^3^ mm^3^), +6cm stimulated 7.87 × 10^4^ mm^3^ (SD = 1.33 × 10^4^ mm^3^), and +8cm stimulated 1.01 × 10^5^ mm^3^ (SD = 1.76 × 10^4^ mm^3^). In sum, the electric field magnitude slightly increased from the baseline CP3-FC3 position to +2 and +4cm, but with a tradeoff of increased volume stimulated (less focal stimulation).

### Round 4: Optimized APPS-tDCS

In a final electrode montage, we optimized tDCS parameters by combining the optimal electrode placement (APPS-tDCS), size (1 × 1cm), and distance (CP3-FC3 +2cm, for an average of 5.12cm apart) that increased the electric field magnitude at the ROI with increased or reasonable focality tradeoff, and compared these results with the original five models (bilateral M1, M1-SO, HD-tDCS, LRPS-tDCS, and APPS-tDCS)(**Figure 7a-c**). Omnibus models were significant for ROI (F(1.7, 348) = 977.2, p < 0.0001), 99^th^ percentile E-field (F(2.6, 512.9) = 898.2, p < 0.0001, and focality (F(2.0, 395.2) = 3079, p < 0.0001.

**Figure 7:**
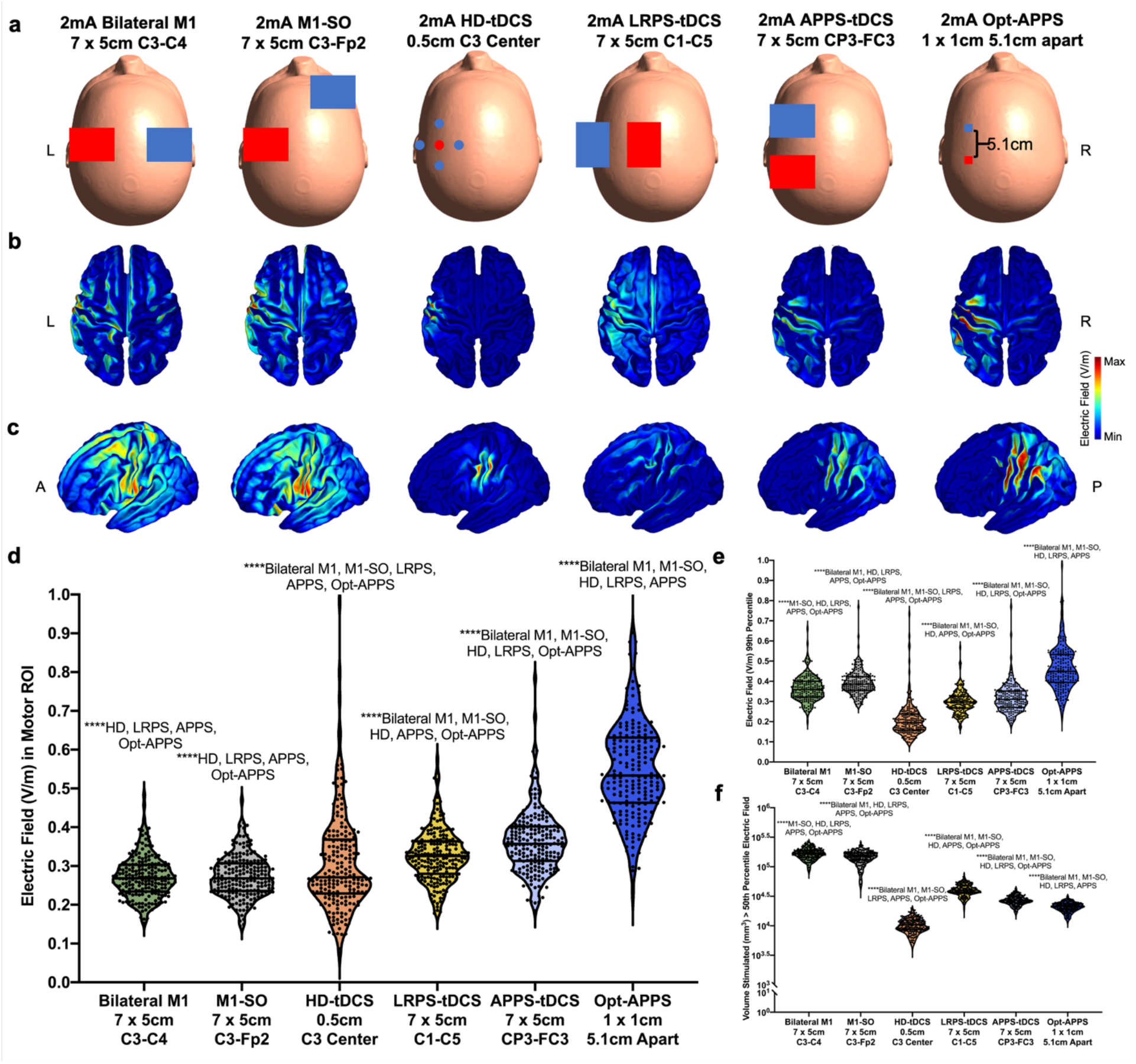
Round 4 Modeling: Optimized APPS-tDCS Produces Double the Electric Field Magnitude at the Same Stimulation Intensity. Combining the principles we learned from Rounds 1-4 modeling, we derived an optimized APPS-tDCS (Opt-APPS) electrode montage. This utilized APPS positioning, with electrode surrounding the cortical motor target in the anterior and posterior directions, 1 × 1cm electrode sizes, and an inter-electrode distance of +2cm (average = 5.12cm inter-electrode distance). Red = anode, blue = cathode. We compared the Opt-APPS placement to the five initial models: bilateral M1, M1-SO, HD-tDCS, LRPS-tDCS, APPS-tDCS (**a**), and observed that Opt-APPS produced qualitatively higher E-fields (**b-c**). **d) EField Magnitude in Motor ROI**. Opt-APPS produced the highest motor ROI E-field of 0.553V/m (****p < 0.0001). **e) E-Field Magnitude Using the 99^th^ Percentile Threshold**. Similarly, Opt-APPS produced the highest global E-field of 0.470V/m. **f) E-Field Focality**. While HD-tDCS still produced the most focal E-field, as measured by the smallest volume equal to or greater than the 50^th^ percentile E-field stimulated, Opt-APPS had more focal E-fields than in every other electrode placement (****p < 0.0001). Thus, Opt-APPS opens the possibility of higher intensity and more efficient stimulation; with the same 2mA stimulation intensity, Opt-APPS produces 4.08mA-like E-fields as it stimulates with 2.04x (more than double) the E-field magnitude and with only 0.12x of the volume stimulated.

Post-hoc Tukey corrected analyses (p < 0.0001) showed that this optimized APPS-tDCS placement produced a significantly higher motor ROI E-field magnitude of 0.554V/m (SD = 0.138V/m) and significantly higher 99^th^ percentile E-field of 0.47V/m (SD = 0.126V/m) than each other condition (**Figure 7d-e; Table 1**). In addition, optimized APPS-tDCS stimulated the significantly lowest volume out of conventional pad electrode montages of 2.14 × 10^4^ mm^3^ (SD = 3.73 × 10^3^ mm^3^)(**Figure 7f; Table 1**).

## Discussion

These data from 3000 E-field models in 200 HCP participants challenge the longstanding assumption that tDCS electrodes should be placed directly over the cortical target to maximally and efficiently stimulate the underlying tissue. Contrary to popular belief and application in thousands of studies, conventionally placing scalp electrodes directly over the intended neural target induces maximal E-fields off-target and midway between the electrodes (**Figure 2a-f**). We leveraged this knowledge into a novel electrode placement called APPS-tDCS, in which we placed the electrodes equidistant and surrounding the cortical M1 target in the anterior and posterior directions. With the same electrode sizes and 2mA stimulation intensity, APPS-tDCS produced a 33% higher E-field with 79% more focal stimulation in the primary motor cortex compared to conventional electrode placements. This group level effect was consistent on an individual level, with 200 of 200 participants experiencing higher E-fields midway between the electrodes than underneath the electrodes (**Figure 3c-d**). Moreover, a similarity analysis between the E-field in the motor target compared to a whole brain 99^th^ percentile threshold E-field revealed that the motor region specific E-field did not contain the top 1% of activated voxels in conventional bilateral M1 and M1-SO electrode placements. However, by using electrodes surrounding the cortical target in LRPS and APPS-tDCS, the maximal Efield was induced at the intended cortical target in each of the 200 participants (**Figure 4a-e**). These three lines of evidence demonstrate that placing tDCS electrodes surrounding a cortical target can produce consistently significantly higher and more focal electric fields than can placing the electrode directly over the target. Thousands of studies have inefficiently stimulated the target with more off-target than on-target effects, which may partially explain the inconsistent behavioral results to date with this inexpensive and noninvasive tool._Using APPS-tDCS, it may be possible to reduce the intersubject variability in experienced E-field(30, 38), or ensure that each participant is more adequately stimulated above a certain cortical E-field threshold, which may result in more consistent behavioral effects(31, 32).

Rounds 2-3 modeling aimed to further optimize tDCS electrode placements and sizes in the novel APPS-tDCS paradigm. These data show that electrode size and distance between electrodes can produce markedly different cortical E-fields from the same 2mA stimulation. When keeping the electrodes an average of 7.12cm apart to match the edge-to-edge distance despite changing electrode sizes, 1 × 1cm electrodes produced the highest motor E-fields of 0.549V/m, compared to 0.428V/m with 7 × 5cm electrodes with a matched average interelectrode distance of 7.12cm (**Figure 5a-f**). We then matched the electrodes for size, using the 7 × 5cm electrodes in each model while moving the electrodes progressively 2cm farther apart. Here we observed that there was a slight increase in E-field magnitude from the original CP3-FC3 electrode positioning to electrodes being farther apart, from 0.363V/m in the original APPS-tDCS positioning to 0.421V/m at +2cm apart, but with decreased focality and more off-target effects with each subsequent move of electrodes farther apart (**Figure 6a-f**).

Taken together, we computed a model using the optimized parameters of the previous four rounds of modeling, positioning the electrodes using Opt-APPS, with 1 × 1cm sizes, and placed an average of 5.12cm apart. This produced an E-field at the motor target of 0.554V/m (**Figure 7a-f**). Compared to conventional electrode montages, this motor cortical E-field translates to a 104% higher (2.04x greater) electric field with 88% less volume stimulated (0.12x spread of stimulation) than in conventional bilateral M1 and M1-SO electrode placements. In other terms, with the same 2mA stimulation input at the scalp, optimized APPS-tDCS produces 4.08mA-like E-fields at the cortical level. Optimized APPS-tDCS also enables more efficient stimulation for typical E-fields (e.g. using 1mA to produce 2.04mA-like E-fields) or higher amounts of cortical stimulation without increasing the scalp dosage. Given an emerging interest in stimulating at higher scalp tDCS intensities(27, 28, 33, 34), perhaps a more effective strategy could be to utilize the principles outlined in this paper to deliver more efficient stimulation at 2mA intensities. Namely, more efficient APPS-tDCS can potentially mitigate adverse effects of higher intensity stimulation including tolerability and focality concerns inherent to higher mA stimulation, although prospective safety and tolerability testing of APPS-tDCS is warranted.

The data we presented in this study have broad scope and inform not only the prospective use of tDCS but also how prior tDCS publications may be interpreted. Whereas there has been a tendency to group in dissimilar electrode positioning or sizes in systematic reviews and meta-analyses, this study points to a need to better understand the effects of these often overlooked variables that can greatly impact the experienced stimulation intensity at the cortical level. This ethos that not all 2mA tDCS is the same resonates with a new meta-analytic approach using retrospective E-field modeling to reinterpret the effects of tDCS on working memory based on variations in electrode positioning and stimulation intensities between studies(39). By better understanding how electrode positioning, size, and distance between electrodes alters the experienced E-field at the cortical target, it may be possible to implement these principles in prospective tDCS studies. Notably, the methods of optimizing tDCS parameters that we outlined in this study do not require different tDCS equipment to be used; rather, researchers can utilize the same tDCS equipment and simply altering the electrode size and positioning can produce profoundly different E-fields at the cortical level. Thus, it is possible to implement APPS-tDCS on a wide scale with minimal alterations to existing practices. Ultimately, we hope that developing a better understanding of how these variables affect Efields could result in more consistent tDCS effects between participants and studies, as tDCS technology has great potential but with inconsistent results to date.

Our finding that the maximum E-field is induced midway between the electrodes additionally has implications for E-field modeling methodology. In particular, many researchers currently measure tDCS E-field magnitude using a whole brain threshold approach, in which they extract the average E-field from the top 1% of activated voxels as a measure of how much stimulation reaches the cortical target (e.g. (40, 41)). While this 99^th^ percentile threshold method appropriately gives a global impression of the tDCS-induced cortical E-fields on the whole brain level, it actually captures the off-target maximum E-field midway between the two electrodes in conventional placements such as bilateral M1 and M1-SO where the anodal electrode is positioned directly over the cortical target. Therefore, if the researcher’s goal is to measure how much stimulation is reaching the intended cortical target, it is best to examine an ROI underneath the anodal electrode or at an anatomical target to determine how much stimulation reaches the target region. In conventional electrode montages, this ROI value is lower than the 99^th^ percentile as the peak E-field effects from placing the electrode directly over the cortical ROI are off-target and between electrodes.

### Future Directions

There are several future directions to build on these large scale E-field modeling data. In addition to APPS-tDCS enabling higher electric fields at the cortical target, there is increasing interest in developing methods of personalizing tDCS dose(30, 32, 38, 41). Researchers have developed E-field methods to individualize tDCS dose such as reverse-calculation modeling(30, 32), which alters tDCS dose at the scalp to produce the same E-field magnitude at a cortical target between participants. A remaining question is whether there is an optimal electric field intensity for personalized reverse-calculation tDCS dosing. Some researchers have proposed to use E-field dosing parameters based on retrospective E-field analyses showing that stimulation above a certain threshold produces greater behavioral effects(31, 32). The Round 1 data in this study could help to refine the optimal electric field intensity for personalized dosing by providing benchmarks for conventional bilateral M1, M1-SO, and HD-tDCS E-fields; these data enable the researcher to make an informed decision whether to stimulate at, below, or above group average E-fields from 2mA stimulation (**Figure 2d-f, Table 1**). These values are also linearly scalable; for instance, to know the average E-fields from 3mA stimulation, the researcher would simply need to multiply the average E-fields from 2mA stimulation by 1.5x. As the tDCS field further refines the best approach for personalizing tDCS dose, we hope that these Round 1 data help to inform feasible E-field dosing parameters.

Furthermore, future research should investigate whether APPS-tDCS at 2mA produces stronger or more consistent behavioral effects as we hypothesize. Moreover, while we focused our 3000 E-field models in the motor cortex as a representative cortical target, more modeling for the prefrontal cortex and other popular stimulation targets could help to inform tDCS parameters and more optimized stimulation strategies in these locations, particularly with the novel APPS-tDCS strategy put forth in this study.

### Limitations

There are several limitations of this study. Electric field modeling is inherently limited by its use of MRI scans to estimate, but not directly measure, how much stimulation reaches the cortex. Important foundational research comparing E-field modeling estimates and actual intracranial recordings established a strong linear relationship between these values (Pearson’s R = 0.88), mitigating some but not all of the concern that E-field modeling is an estimation of cortical stimulation(42, 43). Furthermore, more advanced E-field models, which take different sub-layers of cerebrospinal fluid or skull composition into consideration, may help to further refine the fidelity of E-field models(44, 45). An additional critique is that E-field modeling currently assumes isotropic tissue layers in which the tissue composition is consistent throughout each tissue type, which may not be the case. In one study, researchers demonstrated that anisotropic tissue in deep brain stimulation (DBS) targets could alter electric field modeling estimates by up to 18% compared to homogenous isotropic tissue(46). Further refinements in E-field modeling methodology may further mitigate some of these concerns in the future. Lastly, noninvasive brain stimulation often tests neurophysiology in the motor circuit, as we did in this E-field modeling study, with the hope that motor-induced effects are generalizable to other brain regions such as the prefrontal cortex(47, 48). Prior research has demonstrated that E-fields in the motor cortex are not always fully representative of E-fields in the prefrontal cortex from transcranial magnetic stimulation (TMS)(49). Therefore, while we believe that the effects of tDCS electrode positioning, size, and inter-electrode distance on E-field magnitude and focality will replicate outside of the motor cortex, this needs to be tested forward in the prefrontal cortex and is a topic of ongoing research.

### Conclusions

In summary, we performed 3000 electric field models in 200 HCP participants to test the effects of tDCS electrode positioning, size, and inter-electrode distance on E-field magnitude and focality of stimulation. APPS-tDCS is a method of more optimally stimulating the cortex that can deliver double the on-target electric fields and a fraction of the off-target effects from the same 2mA stimulation intensity as conventional electrode placements. Prospective research using APPS-tDCS could ultimately result in stronger or more consistent transdiagnostic therapeutic effects.

## Methods

### Participants and Scanning Protocol

We accessed the freely available WU-Minn HCP Data Archive (http://www.humanconnectomeproject.org/data/)(35), screening for healthy adult participants with unprocessed T1w and T2w structural MRI scans. Of the 1113 individuals with deidentified structural scans, we randomly selected 200 (100 male/100 female). As part of the deidentification process for the public database, only general age ranges are reported in the HCP data. As such, we chose to represent the age ranges available in the dataset to the best of our ability-33F/33M in the 22-25 range, 34F/34M in the 26-30 range, and 33F/33M in the 31-35 range.

Structural MRI scans in the HCP database were acquired with Siemens MAGNETOM 3T scanners with 32-channel head coils. T1w scan parameters were as follows: TR = 2400ms, TE = 2.14ms, flip angle = 8°, field of view = 224mm x 224mm x 180mm, voxel size = 0.7mm^3^. T2w scans were acquired at the following parameters: TR = 3200ms, TE = 565ms, field of view = 224mm x 224mm x 180mm, voxel size = 0.7mm^3^. Full imaging parameters are available at the HCP database, *Appendix I: HCP scan protocols*.

### Electric Field Modeling and Analysis

For segmentation and meshing of T1w and T2w structural MRI scans, we used headreco (https://simnibs.github.io/simnibs/build/html/documentation/command_line/headreco.html)(50). Headreco utilizes SPM12 (https://www.fil.ion.ucl.ac.uk/spm/) and CAT12 (http://www.neuro.uni-jena.de/cat/) to segment each scan into skin, skull, cerebrospinal fluid, white matter, grey matter, and eyes, and generate a finite element model (FEM) containing each tissue type (**Figure 8**). Following segmentation, headreco meshed the tissue layers together. We visually evaluated the integrity of the FEM mesh to ensure that the tissue segmentations were clearly delineated between layers. We assigned default tissue conductivity values to each tissue layer: skin: 0.465S/m, bone: 0.01S/m, cerebrospinal fluid: 1.654S/m, grey matter: 0.275S/m, white matter: 0.126S/m, and eyeballs: 0.50S/m, and with a mesh density of 0.5 nodes per mm^2^. For Efield modeling, we used SimNIBS 3.2.3 (https://simnibs.github.io/simnibs/build/html/index.html)(51). Electrode positions, sizes, and locations for each model are indicated in **Table 1** and **Figures 2 and 4–7** summarizing Rounds 1-4 of modeling.

**Figure 8:**
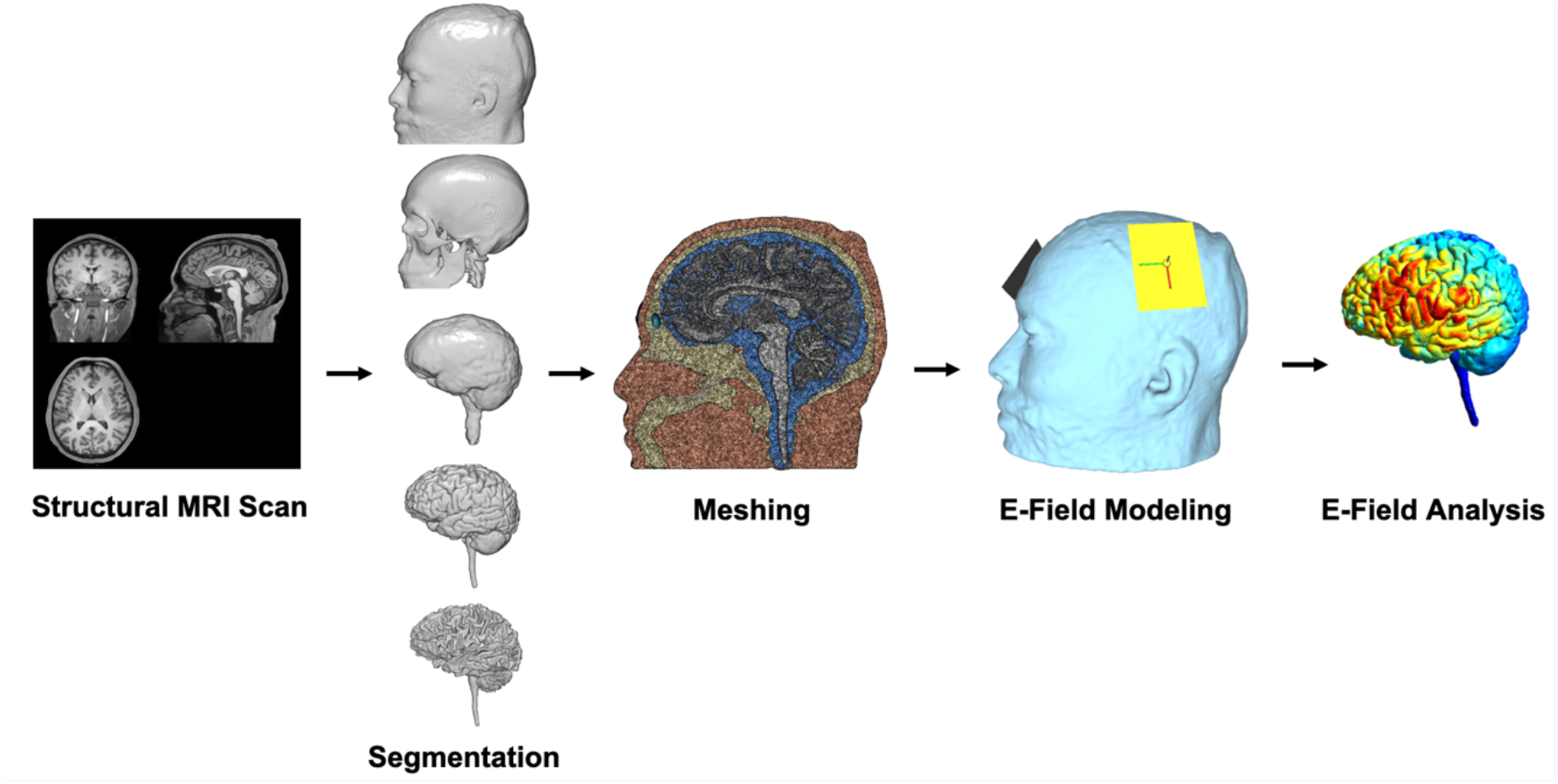
E-Field Modeling Pipeline. This figure shows the five main steps of E-field modeling. First, the researcher acquires structural MRI scans (T1w and T2w). Next, the structural MRI scan is segmented into different components (top to bottom: skin, bone, cerebrospinal fluid, grey matter, and white matter). Third, the tissue segmentations are combined into a 3D volumetric mesh. Fourth, E-field modeling uses experimentally determined tissue conductivity values to simulate how much electric current applied at the scalp reaches the cortex. Displayed is an example in which 7 × 5cm electrodes were placed over the left primary motor cortex (M1) and right supraorbital (SO) cortex. Lastly, the researcher performs E-field analyses. See **Figure 1** for detailed description of E-field analyses in this study, which included a region of interest (ROI) analysis, 99^th^ percentile threshold analysis, and focality analysis as a measure of volume stimulated at or above the 50^th^ percentile E-field.

For each model, we computed three types of data (**Figure 1**). First, we extracted the average E-field in a 10mm radius spherical region of interest (ROI) that was centered at the grey matter voxel directly underneath the center of the projection from C3 at the scalp level. This motor ROI was individually placed for each person and was the same for each model regardless of the tDCS electrode positioning. Second, we examined the E-field in the top 1% of voxels by using a whole brain 99^th^ percentile threshold. Third, we measured focality by summating the volume of voxels stimulated above the whole brain 50^th^ percentile electric field. The greater the volume stimulated, the more diffuse and less focal the stimulation. In each model, we extracted the normal component of the E-field (normE).

### Statistical Analysis

Significance was set at p < 0.05 for all statistical analyses (two-tailed). Rounds 1-4 of modeling were analyzed using repeated measures ANOVAs in GraphPad PRISM 9.0.1. All post-hoc analyses were Tukey corrected with significance level set at p < 0.05. Intraclass correlation coefficients were calculated using IBM SPSS Version 25.0.

### Data Availability

WU-Minn HCP data are freely available online (http://www.humanconnectomeproject.org/data/). Requests for segmented and meshed scans, E-field models, or further documentation are available upon reasonable request.

## Author Contribution Statement

KAC conceived of the study, performed all data analyses, and wrote the initial manuscript draft. MSG edited the initial manuscript and provided feedback.

## Acknowledgements

Data were provided [in part] by the Human Connectome Project, WU-Minn Consortium (Principal Investigators: David Van Essen and Kamil Ugurbil; 1U54MH091657) funded by the 16 NIH Institutes and Centers that support the NIH Blueprint for Neuroscience Research; and by the McDonnell Center for Systems Neuroscience at Washington University. Funding was also provided by the National Center for Neuromodulation and Rehabilitation (NC NM4R) (Principal Investigator: Steven A. Kautz; 2P2CHD086844) and the Center of Biomedical Research Excellence (COBRE) in Stroke (Principal Investigator: Steven A. Kautz; 5P20GM109040). The Brain Stimulation Laboratory (Principal Investigator: Mark S. George) also receives funding from the Tiny Blue Dot Foundation.

## Supplementary Section 1

This supplementary section accompanying Caulfield & George describes some of our tDCS electric field (E-field) modeling in 200 Human Connectome Project (HCP) participants. This round of modeling (Round 1.5), which we computed after Round 1 and prior to Round 2, tested the effects of electrode size and distance in anterior posterior pad surround (APPS)-tDCS by placing electrodes of differing sizes (7 × 5cm, 5 × 5cm, 3 × 3cm, and 1 × 1cm) centered at individual CP3-FC3 coordinates for each participant (**Supplementary Figure 1a**). A caveat of this approach was that the inter-electrode distance was larger with smaller electrodes centered at the same locations. The 1 × 1cm electrodes had the largest average inter-electrode distance of 7.12cm, compared to 5.12cm for the 3 × 3cm electrodes and 7.12cm for the 5 × 5cm and 7 × 5cm electrodes. Thus, differences between these models pointed to an effect of electrode size and/or distance, but with no way to identify which variable caused this effect or how much each variable contributed to the observed effects.

### Round 1.5: Effect of Electrode Size and Distance in APPS-tDCS

Having established that placing electrodes surrounding the cortical motor produces higher and more focal E-fields in Round 1 modeling, examining the ROI E-field underneath vs. between electrodes, and in ICC similarity analyses, we next tested variables that might further affect the E-fields delivered to the cortex using the APPS-tDCS montage. In this Round 1.5 of modeling, we compared the effects of placing electrodes of differing sizes (7 × 5cm, 5 × 5cm, 3 × 3cm, and 1 × 1cm) at the CP3-FC3 coordinates in the original APPS-tDCS model on the same ROI, 99^th^ percentile, and focality measures in 200 models per condition (1 per person, paired between models)(**Supplementary Figure 1a-c**).

Qualitatively and quantitatively, electrode size altered E-fields at the motor ROI target (F(1.1, 228.3) = 1700, p < 0.0001 and with the 99^th^ percentile metric (F(1.1, 217) = 1112, p < 0.0001 (**Supplementary Figure 1d-e**). Post-hoc Tukey corrected comparisons revealed that each electrode montage for the ROI and 99^th^ percentile measures significantly differed at the p < 0.0001 significance level for each comparison. For the ROI measure, the 1 × 1cm APPS-tDCS electrodes had the highest E-field of 0.549V/m (SD = 0.133V/m), followed by 3 × 3cm with 0.512V/m, SD = 0.122V/m, 5 × 5cm with 0.416V/m (SD = 0.095V/m), and the original 7 × 5cm montage with an E-field of 0.363V/m (SD = 0.081V/m). For the 99^th^ percentile measure, the 1 × 1cm APPS-tDCS electrodes also produced the highest E-field of 0.489V/m (SD = 0.135V/m), which was significantly higher than the E-field produced by 3 × 3cm electrodes of 0.444V/m (SD = 0.107V/m), 5 × 5cm electrodes of 0.347V/m (SD = 0.075V/m), and the 7 × 5cm electrodes of 0.316V/m (SD = 0.069V/m).

For focality, the different electrode montages produced relatively the same amount of spread of electric fields. The F-test was statistically significant, F(1.3, 255.8) = 47.4, p < 0.0001. The 5 × 5cm APPS-tDCS electrode placement produced the most focal electric fields of 2.49 × 10^4^ mm^3^ (SD = 4.54 × 10^3^ mm^3^), compared to 2.66 × 10^4^ mm^3^ (SD = 4.68 × 10^3^ mm^3^) for 1 × 1cm electrodes, 2.67 × 10^4^ mm^3^ (SD = 4.33 × 10^4^ mm^3^) for 3 × 3cm electrodes, and 2.75 × 10^4^ mm^3^ (SD = 4.98 × 10^3^ mm^3^) for 7 × 5cm electrodes (**Supplementary Figure 1f**).

**Supplementary Figure 1:**
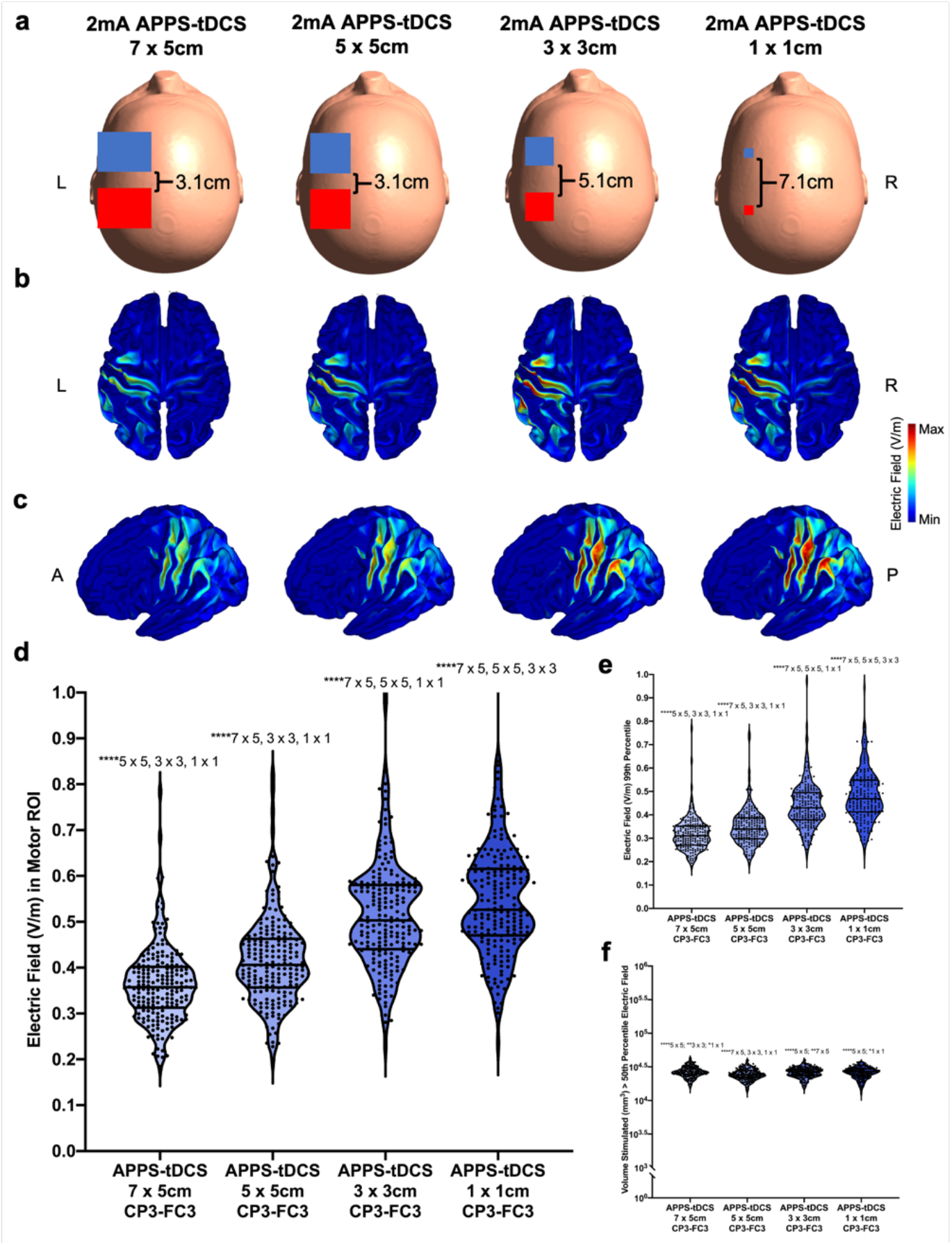
Round 1.5 Modeling: tDCS Electrode Size and Position Significantly Affect E-Field Magnitude and Focality. In Round 1.5 of modeling, we positioned electrodes of 7 × 5cm, 5 × 5cm, 3 × 3cm, and 1 × 1cm at the CP3-FC3 EEG locations. Red = anode, blue = cathode. **a) Electrode Placements in Each Montage. b-c) E-Fields from 200 Models Projected Into fsaverage Space for Axial (b) and Sagittal (c) Orientations**. In each projection, 200 individual models were averaged into fsaverage space to visualize the E-field magnitude and spread for each electrode orientation. **d) E-Field Magnitude in Motor ROI**. Electrode size significantly impacted the E-field within the motor cortical target (****p < 0.0001). When placed at the CP3-FC3 locations, 1 × 1cm electrodes produced a mean ROI E-field of 0.549V/m, significantly greater than the ROI E-fields of 3 × 3cm (mean = 0.518V/m), 5 × 5cm (mean = 0.416V/m), and 7 × 5cm (0.363V/m). **e) E-Field Magnitude Using the 99^th^ Percentile Threshold**. 99^th^ percentile E-fields were congruent with the ROI E-fields, such that the 1 × 1cm 99^th^ percentile E-field was significantly greater than the E-field from each other electrode placement. **f) E-Field Focality**. The volume stimulated differed between electrode sizes, with 5 × 5cm positioning producing the most focal E-fields (****p < 0.0001, **p < 0.01, *p < 0.05). A caveat of Round 2 modeling is that by placing electrodes of different sizes and centering them at CP3-FC3, the inter-electrode distance differed between an average of 7.12cm (1 × 1cm) and 3.12cm (7 × 5cm and 5 × 5cm). Rounds 3-4 of modeling aimed to dissociate Round 2’s conflated variables of electrode size and inter-electrode distance to isolate these effects.

